# Genomic and functional insights on *Priestia megaterium* MOD5IV: Enhancing Metal Phytoremediation Potential in Arid Environments

**DOI:** 10.1101/2025.04.15.648928

**Authors:** Luis Pouchucq, Andrés E. Marcoleta, Cristian Becerra, Carola Bahamondes, Pablo Lobos-Ruiz

## Abstract

**Aims:** Metal contamination poses a global threat due to its widespread occurrence and the high toxicity of these elements. Phytoremediation has emerged as a preferred approach for the bioremediation of metal-contaminated soils. The search for microorganisms facilitating phytoremediation, especially plant-growth-promoting bacteria (PGPB), has become critical to advance ecosystem remediation efforts. This research aimed to characterize in-depth a *Priestia megaterium* strain isolated from multimetal contaminated soils located at the Atacama Desert, showing potential for bacteria-assisted phytoremediation.

**Methods and Results:** The strain MOD5IV exhibited notable PGPB features: phosphate and potassium solubilization, nitrogen fixation, phytohormone production, and growth promotion of *Arabidopsis thaliana*. Genomic analysis revealed a 5,254,635 bp chromosome plus nine plasmids, hosting over 6,000 genes. Functional annotation identified genes associated with resistance to copper, cadmium, lead, mercury, zinc, and cobalt. Also, genes linked to PGPB capabilities as siderophore-production, nutrient-solubilization, IAA-synthesis, and nitrogen-fixation. Accordingly, MOD5IV exhibited robust tolerance to multiple metals and enhanced the phytoremediation potential of *Caesalpinia Spinosa* (Mol.) in laboratory trials.

**Conclusions:** MOD5IV proved to have promising traits for microbe-assisted phytoremediation of metal-contaminated soils.

**Impact Statement:** This study contributed to the characterization of new native multi-metal-resistant PGPR bacteria for phytoremediation of metal-contaminated soils. Increasing the evidence of the Atacama Desert as a source of microbiological solutions for climate adaptation and environmental remediation.

## INTRODUCTION

Soil contamination by heavy metals is a significant concern for governments worldwide. Various regulations establish concentration limits for several heavy metals based on land use (Sánchez-Castro 2023, Parviainen 2022). Additionally, regulations require polluting companies, such as those in the mining industry, to take actions to prevent pollution or to remediate contaminated sites. Governments are making concerted efforts to address soil contamination by heavy metals and mitigate its effects on human health and ecosystems (Liu 2021). There are various strategies for the remediation of these contaminated soils, most of which are based on physical or mechanical methods such as excavation and landfilling, surface capping, chemical solubilization and stabilization, and soil washing, among others. Furthermore, biological strategies are generally based on microorganism-mediated or plant-mediated bioremediation methodologies (Souza 2020, Azhar 2022). The best efficiencies at the field scale have been reported combining physical and biological strategies (Sánchez-Castro 2023).

Among the biological strategies, plant-mediated bioremediation, called phytoremediation, is a widely accepted approach due to its economic feasibility and eco-friendliness (Jacob 2018). However, many factors, such as high metal concentrations, nutrient deficiency, soil structure, and physicochemical factors like pH and salinity, limit biomass production and, therefore, the phytoremediation efficiency of plants (Alves 2022). Improving the phytoremediation efficiency of plants at higher metal concentrations is a promising approach, that may be achieved by utilizing plant growth-promoting bacteria (PGPB) (Manoj 2020; Simmer 2022).

PGPB comprise species from a broad number of genera, including free-living bacteria, those that establish specific symbiotic relationships with plants, bacterial endophytes that colonize plant interior tissues, and cyanobacteria (blue-green algae) (Alves 2022). These bacteria can fix nitrogen, solubilize phosphate and potassium, and produce phytohormones, significantly reducing the need for chemical fertilizers and modulating the effects of environmental stress and pathogen control (Glick 2012, Romero-Estonllo 2023, Asad 2018, Sijilmassi 2020, Khoshru 2020, Backer 2018, Nazari 2020, Myo 2019, Hernández-Soberano 2020, Naqqash 2020). Several bacterial strains isolated from the plant rhizosphere have been reported to improve the productivity and health of plants under metal contamination stress, making them a significant component in phytoremediation processes (Kamaruzzaman 2020, Srivastava 2013, Xu 2018, Wu 2019, Liu 2022).

Among them, *Priestia megaterium* (formerly *Bacillus megaterium*) strains have demonstrated notable plant growth-promoting (PGP) properties, including potassium solubilization and nitrate reduction (Wu 2023, Wang 2020). These strains also exhibit bioremediation capabilities for environments contaminated with heavy metals like zinc (Zn) and cadmium (Cd) (Bhatt 2020, Wu 2019) and can induce plant resistance to environmental stresses or pathogens (Cui 2023, Wang 2020, Chakraborty 2006, Zhou 2016).

In this study, we report on the functional and genomic characterization of the PGP and multi-metal-resistant strain *P. megaterium* MOD5IV, isolated from the rhizosphere of *Caesalpinia spinosa* (Mol.) plants thriving in a former mine tailing deposit in the semiarid edge of Atacama Desert. Additionally, we investigated its capability to promote growth and enhance the phytoremediation performance.

## MATERIALS AND METHODS

### Strain isolation

Strains were isolated from the rhizosphere of *Caesalpinia spinosa* plants growing on former tailings from artisanal copper mining near Andacollo, a traditional mining city located in Elqui Province in northern Chile (30°15.381’S 71°04.551’W). Soil samples (1 g) collected from the mine tailings were added to 10 mL of sterile water and vortexed vigorously for 1 min. Serial tenfold dilutions were prepared and then spread on Soy Trypticase Agar (TSA) medium (Difco®). Plates were incubated at 28°C for 24 h.

### Basic qualitative PGPB analysis

Nitrogen fixation was determined by growth on Ashby-agar medium (0.2 g/L KH_2_PO_4_, 0.2 g/L MgSO_4_, 0.2 g/L NaCl, 0.2 g/L CaSO_4_, 5.0 g/L CaCO_3_, 5.0 g/L sucrose, 5.0 g/L glucose, and 15 g/L agar) (Becking 2006). Phosphate and potassium solubilization index were determined cording Sijilmassi (2020), using Pikovskaya agar medium (10.0 g/L glucose, 0.2 g/L NaCl, 0.002 g/L FeSO_4_, 0.002 g/L MnSO_4_, 0.2 g/L KCl, 0.1 g/L MgSO_4_, 0.5 g/L (NH_4_)_2_SO_4_, 0.5 g/L yeast extract, 5.0 g/L Ca_3_(PO4)_2_, and 15 g/L agar) and modified Pikovskaya agar medium (10.0 g/L glucose, 0.2 g/L NaCl, 0.002 g/L FeSO_4_, 0.002 g/L MnSO_4_, 0.2 g/L KCl, 0.1 g/L MgSO_4_, 0.5 g/L (NH_4_)_2_SO_4_, 0.5 g/L yeast extract, 5.0 g/L KNO_3_, and 15 g/L agar), respectively. Siderophore production was tested using the chrome azurol S method in solid medium (CAS-agar), as described by Alexander and Zuberer (Alexander 1991). The bacterial cultures were incubated at 28°C for 24 h. Orange halos around the colonies indicated siderophore production.

IAA production was determined colorimetrically, according to Glickmann (1995). The isolates were inoculated in Soy Trypticase Broth (TSB) medium with and without tryptophan (500 μg/ml) and incubated at 28°C for 4 days with shaking at 250 rpm. A 1 mL aliquot was removed from each tube and centrifuged at 10,000 rpm for 10 min. 200 μL of the supernatant from each sample was transferred to a 96-well microplate, to which 50 μL of Salkowski reagent (12 g/L FeCl_3_ and 7.9 M H_2_SO_4_) were added. The mixture was incubated at room temperature for 30 min, and the absorbance was measured at 530 nm using a microplate spectrophotometer (EPOCH-II BioTek). The IAA concentration in the culture was determined using a calibration curve of pure IAA (Supelco®) as a standard, following linear regression analysis.

### Phosphate uptake quantification

PO₄ was quantitatively determined using the Mo-Blue Method (Watanabe 1965, Nautiyal 1999). A pre-culture was incubated in 10 mL of TSA medium overnight at 29°C with agitation (150 rpm). The following day, the culture was adjusted to 10^8 CFU/mL, and 100 μL were inoculated into 10 mL of PKV medium. The cultures in PKV medium were incubated at 29°C with constant agitation (150 rpm) for 72 h. One milliliter of the culture was clarified by centrifugation (12,000 x g), and 250 μL of the supernatant were placed in a microcentrifuge tube and digested by adding concentrated sulfuric acid in a 1:1 v/v ratio. 200 μL of the digested samples were placed in triplicate in a 96-well microplate. 100 μL of Mo-Blue reagent was added to each sample and mixed gently by pipetting. Absorbance was recorded using a microplate spectrophotometer (EPOCH-II BioTek) set at 820 nm. A calibration curve was constructed with a PO₄ standard (KH₂PO₄, Certipur®, Supelco).

### Arabidopsis thaliana growth promotion

*A. thaliana* (Col-0) seeds were treated with 2% Tween 20 for 10 min with gentle agitation. The seeds were then rinsed thrice with sterile distilled water. Afterward, the seeds were soaked in 70% ethanol for 30 s and rinsed three times with sterile distilled water. Subsequently, the seeds were dipped in 1% sodium hypochlorite for 10 min and finally rinsed with sterile distilled water five times. Seeding was carried out in polycarbonate Petri plates with Murashige & Skoog (MS) medium supplemented with sucrose (10 g/L) and agar (8 g/L) at pH 5.7. Three seeds were placed into each plate at 2 cm from each other and incubated at 4°C for 48 h for stratification. After the stratification period, the plates were placed vertically (at a 65° angle) in an incubator with a photoperiod of 16:8 h (light:dark) at 24°C for 6 days. On the sixth day of culture, 20 μL of MOD5IV at 10^8 CFU/mL were inoculated 2 cm from the seedling roots. The seedlings were incubated for an additional 8 days under the same conditions. Finally, the length of the primary root, number of secondary roots, rosette diameter, leaf area, and number of true leaves were measured. This assay was carried out in triplicate, with a total of 9 plants for each isolate.

### IAA quantification by HPLC

MOD5IV (0.3 mL of 10^8 CFU/mL culture) was inoculated into 30 mL of TSB medium, with and without tryptophan (500 μg/mL), in 50 mL centrifuge tubes, and incubated at 28°C for 72 h with shaking at 250 rpm. After incubation, the culture was centrifuged at 7,000 rpm for 10 min for cell clarification. 25 mL of the supernatant were adjusted to pH 2.5 with HCl. IAA was extracted with ethyl acetate in a 1:3 ratio, vortexing for 1 min. The ethyl acetate was evaporated at 40°C overnight, and the resultant powder was dissolved in 3 mL of methanol. The sample was analyzed by HPLC (Ultimate 3000 Thermo®) using a pre-equilibrated reverse-phase column (C18 LiChrosphere 5 μm, Merck®) under isocratic conditions with a mobile phase of water:acetonitrile: acetic acid (85:15:1). The chromatograms were run at a flow rate of 1 mL/min at 20°C. Detection was performed by UV light absorption at 252 nm. A calibration curve was constructed using an HPLC-grade standard of IAA (Supelco®) (Shao 2015, Tien 1979).

### Quantification of ammonia production

Nitrogen fixation was determined based on ammonia production using the Nessler’s method (Kamaruzzaman 2020, Wu 2019). MOD5IV preinoculum was prepared in TSB and incubated overnight at 29°C with shaking at 150 rpm. The next day, CFUs were determined using the microdroplet technique. Inocula were prepared in triplicate tubes with 20 mL of peptone water (20 g/L peptone and 30 g/L NaCl). The tubes were incubated at 29°C for 5 days with shaking at 150 rpm. After incubation, CFU was determined using the microdroplet technique. Bacterial cells were eliminated by centrifugation at 12,000 rpm for 10 min. Then, 200 µL of the culture supernatant was transferred to new microcentrifuge tubes. Subsequently, 1 mL of Nessler’s reagent was added. The optical density was measured at 450 nm using a sterile 96-well microplate. A calibration curve was generated using an ammonium standard (NH_4_Cl Certipur® Supelco) on the same plate.

### Metal exposure and MIC determination

The strain was inoculated from a previous culture adjusted at 10^8 CFU/mL in a proportion of 1:100 in TSB medium supplemented with previously diluted gradient of metal concentrations: copper (CuSO_4_), mercury (HgCl_2_), chromium (K_2_Cr_2_O_7_), lead (Pb(C_2_H_3_O_2_)_2_), cadmium (CdCl_2_) and arsenic (As_2_O_3_) (ReagentPlus®, Sigma-Aldrich). pH was adjusted to 7.5. The cultures were placed in triplicate into a sterile 96-well microplate and incubated at 28°C with agitation of 200 rpm into an Epoch-II (BioTeck) microplate reader. Cell proliferation was determined by absorbance at 600 nm, measuring every 0.5 h for 24 h. The minimum inhibitory concentration (MIC) was determined for each metal as the concentration needed to avoid culture growth in the conditions previously mentioned.

### Promotion of *C. spinosa* growth in metal-contaminated substrate

*C. spinosa* seeds were treated with concentrated sulfuric acid for 45 min. The solution was then neutralized with sodium bicarbonate. The seeds were rinsed with distilled water and placed into plastic containers containing 80 g of contaminated soil from the same mine tailing where the MOD5IV strain was collected, amended in a 50:50 ratio with sterilized plant substrate based on Sphagnum sp. H2–H4 von Post (Kekkila® Professional substrate). The MOD5IV inoculum was grown overnight in TSB medium, and the cell density was adjusted to 10^8 CFU/mL. One milliliter of this inoculum was added to each substrate container. Control containers received TSB medium without bacteria. One seed was placed in each container, and the containers were incubated at 28°C for 60 days with a 16:8 light-dark photoperiod. Five replicates were prepared for each condition.

After the culture period, seedlings were carefully extracted and rinsed with distilled water. Growth parameters, including root length and dry mass, were recorded. The dried tissues were used to determine copper (Cu) concentration. The plant material was calcined in a muffle furnace until white ash was obtained. The white ash was suspended in nitric acid and diluted with distilled water (1:10). The solutions were clarified by filtering, and Cu concentrations were quantified by atomic absorption spectroscopy using an AAnalyst™ 700, PerkinElmer instrument. Metal concentrations in the substrate were quantified by X-ray fluorescence (XRF). A soil (tailing) sample was collected during bacterial isolation, from a 1 m × 1 m area to a depth of 15 cm using a shovel. The amount of soil was then reduced using the quartering method (ASTM International 2018) to a representative 1 kg sample. The tailing material, both alone and amended, was dried in an oven at 60°C until no mass loss was observed. The samples were analyzed using X-ray fluorescence (XRF) with an S1 Titan 600 Handheld XRF Analyzer (Bruker).

### Genomic sequencing, assembly and annotation

Genomic DNA was purified from an overnight culture of the MOD5IV strain grown in TSA, using the genomic extraction kit Wizard (Promega®) following the manufacturer’s recommendations. Genomic DNA was sequenced by Illumina® and Nanopore® platforms. Illumina sequencing (100-pb paired-end reads) was performed through the TruSeq Nano DNA Kit for library preparation and run in a XLEAP-SBS machine (Illumina SBS technology). The obtained reads (15,374,518) were quality-checked by the FastQC 0.11.9 software (FastQC, 2025). The Nanopore sequencing library was prepared using the Ligation Sequencing kit LSK-110, following the manufacturer’s guidelines. 5–50 fmol of the library were loaded into a FLO-MIN106D flow cell and sequenced using a MinION device (Nanopore Technologies Ltd., Oxford, UK). The quality and length of the Nanopore reads were assessed by Nanoplot from the Nanopack toolkit (Oxford Nanopore). Unicycler 0.4.9b (Wick 2017) was used in hybrid mode for genome assembly. Gene annotation was performed using the Rapid Annotation program by the Subsystem Technology (RAST) server (Overbeek 2014). Additional functional annotation was carried out using the KEGG (https://www.kegg.jp/kegg/kegg2.html) database. The Resistance Gene Identifier (RGI) and the CARD database (https://card.mcmaster.ca/analyze/rgi) (Alcock 2020) were used for resistome prediction.

## RESULTS

### Isolation and initial characterization

To screen for the presence of metal-resistant bacteria with high-efficiency plant growth promotion (PGP) properties, the rhizosphere soil of *Caesalpinia spinosa* was selected because it was thriving on a former mine tailing deposit with high concentrations of metals (Supplementary Table 1). The isolation of pure strains was performed on copiotrophic TSA medium agar plates. A series of bacterial strains were obtained and screened for PGP properties, including nitrogen fixation, phosphate and potassium solubilization, IAA production, and promotion of *Arabidopsis thaliana* growth. The bacterial isolates that most effectively induced the growth of *A. thaliana* are shown in Figure 1A. Of these, the MOD5IV isolate exhibited the greatest induction of primary root growth.

**Figure 1.**
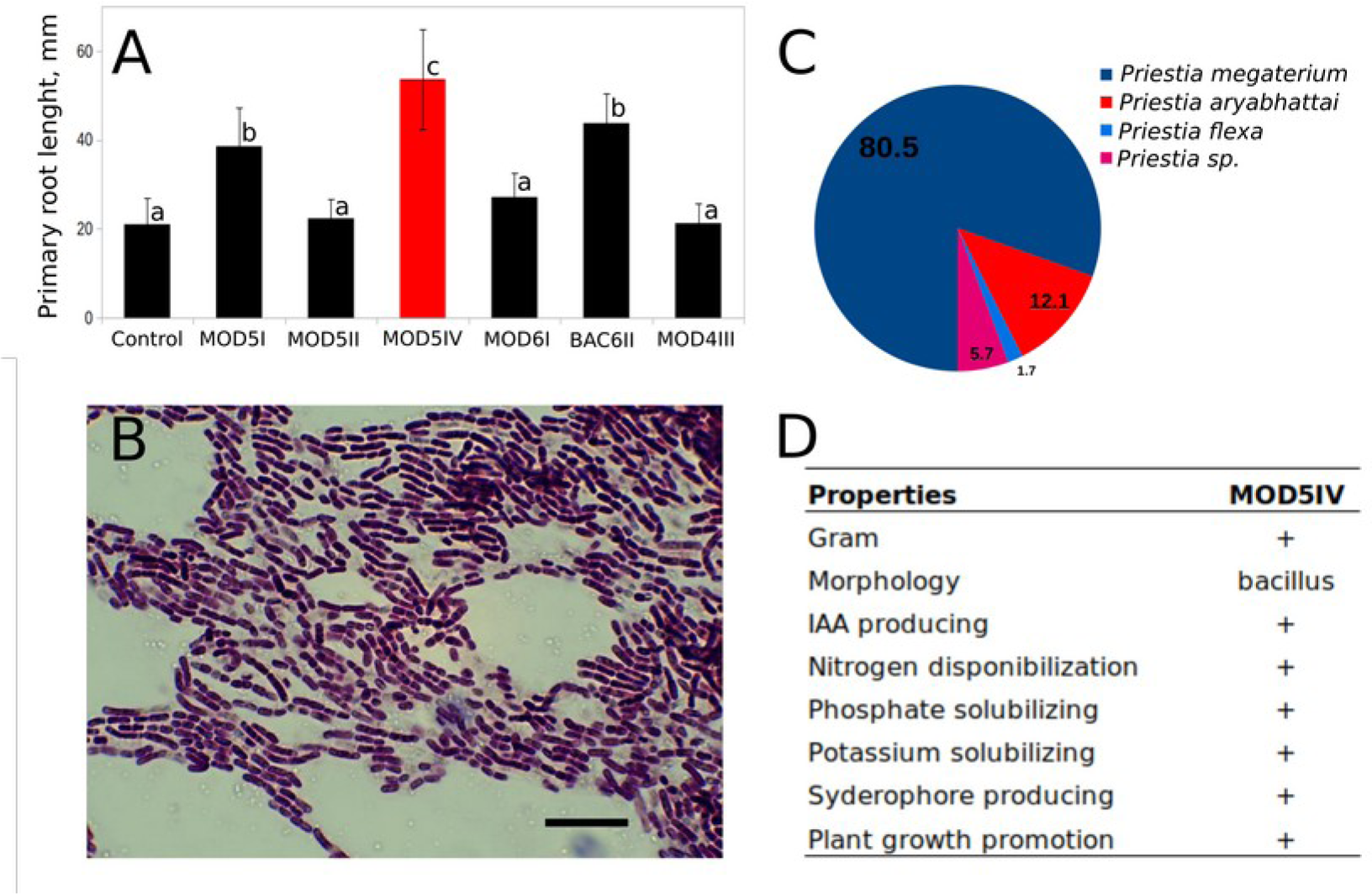
Selection of Isolates, Plant Growth Promotion Capacity, and General Characterization of Strain MOD5IV. A) Arabidopsis thaliana primary root growth promotion by the isolated strains. The red column represents the strain MOD5IV, which had the highest plant growth promotion capacity. a, b and c indicate significant differences estimated using Two-way ANOVA (p < 0.05). Parametricity was estimated using the Kolmogorov-Smirnov Test of Normality. N = 9. B) MOD5IV morphology as revealed by bright-field microscopy and Gram staining (scale bar indicates 25 µm). C) Species distribution map of MOD5IV 16S rDNA compared to the NR database (numbers represent the percentage of representation of the 100 best hits in BLAST analysis). D) Summary of the morphological characteristics and plant growth promotion properties of strain MOD5IV.

The MOD5IV isolate formed white colonies with lobulated margins on TSA medium and showed a bacillar Gram-positive morphology under microscopic analysis (Figure 1B), with a cell length of 10.5 ± 2.4 μm. To confirm the taxonomic position of MOD5IV, we sequenced the 16S rDNA (Accession number MN744240.1). BLAST analysis indicated the highest similarity with *Priestia megaterium* SR01-1 and *Priestia megaterium* DK2 (100% identity for both). The species distribution map of the 16S rDNA gene compared to the 16S rRNA database is shown in Figure 1C, where more than 80% of the hits obtained corresponded to sequences annotated as belonging to the taxa *Priestia megaterium* (NCBI: txid1404). This evidence, combined with the comparison results of 16S rDNA gene sequence-based and whole-genome sequence-based phylogenetic trees elaborated with TYGS (Meier-Kolthoff 2022) (Supplementary Figure 1), allowed us to conclude that MOD5IV belongs to the *Priestia megaterium* species. This represents the first reported complete genome sequence of the taxa (*P. megaterium*) for the arid zones of South America (Adeniji 2025).

### MOD5IV plant growth promotion properties

The basic plant-growth-promoting properties of MOD5IV were examined (Figure 1D). MOD5IV was able to solubilize phosphate and potassium in solid PKV agar medium. The solubilization indexes for both nutrients were 8.6 and 35.3, respectively, after 72 h of culture. PO₄ solubilization was also quantified using the Mo-Blue method, determining that MOD5IV was capable of incorporating 69 ± 8 µg(PO₄)/mL directly from inorganic Ca₃(PO₄)₂ present in the PKV medium during 72 h of culture. Similarly, MOD5IV was able to grow in nitrogen-deficient Ashby medium, indicating the presence of a nitrogen fixation mechanism. Quantification of NH₄ production through the Nessler reaction indicated that MOD5IV produced 100.6 ± 3.6 µgNH4/mL in a 5-day culture. Siderophore production was evidenced by the formation of a characteristic halo on CAS-agar medium. The production of the phytohormone IAA was assessed colorimetrically using the Salkowski method and also by HPLC analysis. In a 72-h culture, the MOD5IV strain produced 40.9 μg/mL of IAA, according to the Salkowski method, and 0.3 μg/mL of IAA using HPLC analysis. This difference of two orders of magnitude between the methods is due to the Salkowski method reacts positively with various auxins and their precursors. In contrast, the HPLC method can differentiate between different types of auxins and precursors as indole-3-lactic acid (ILA), tryptamine, indole-pyruvic acid, among others (Guardado-Fierros 2024, Tien 1979).

The plant growth-promoting properties (PGPR) of MOD5IV were tested using a co-culture method with *Arabidopsis thaliana* in solid plant MS medium. Results shown in Figure 2 indicate that the presence of MOD5IV induced a significant increase in all of the growth parameters tested: length of the primary root (Figure 2A), number of secondary roots (Figure 2B), and rosette diameter (Figure 2C). The increase was even more pronounced in the parameters related to the root. Representative images are shown in Figure 2D.

**Figure 2.**
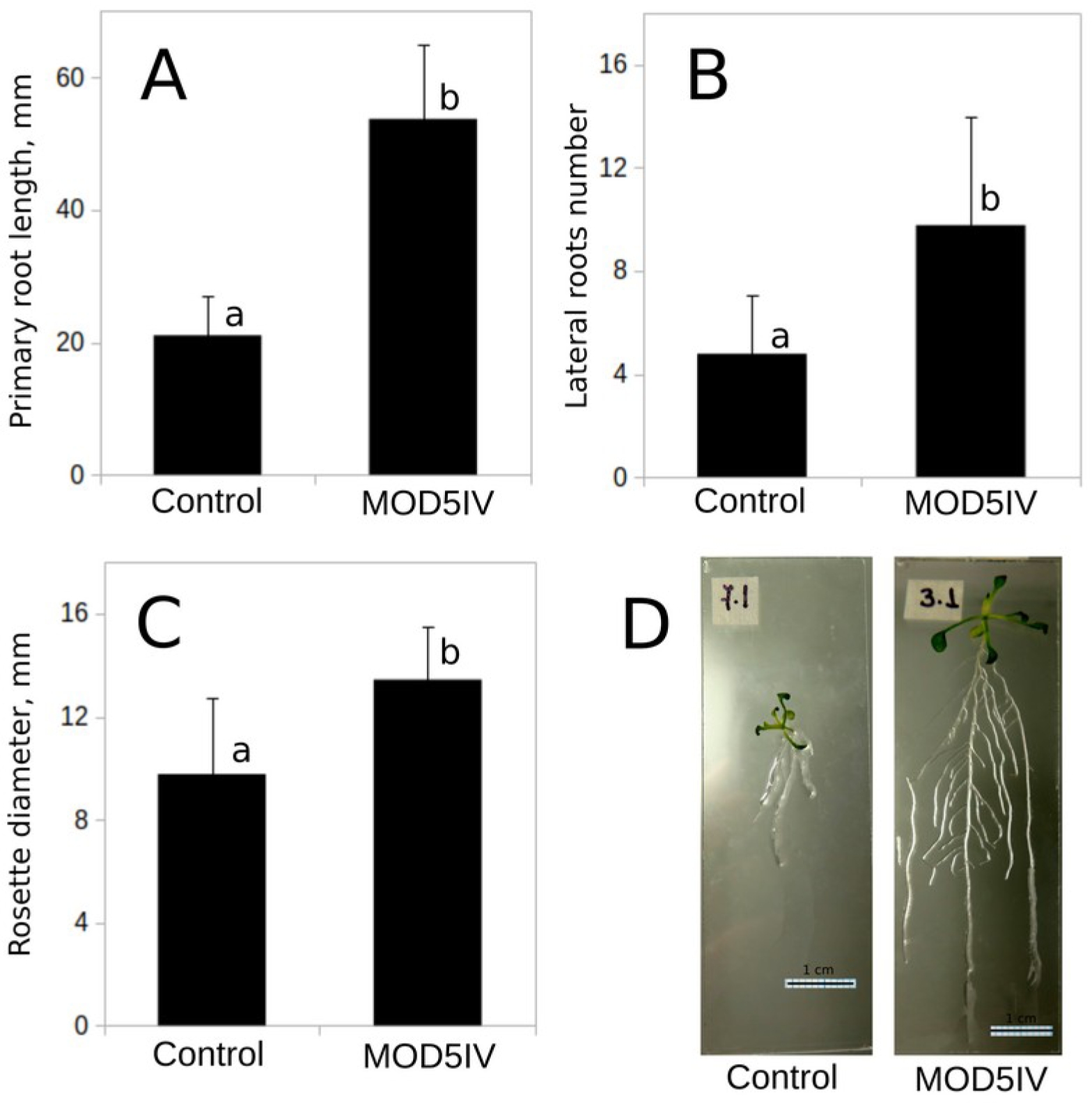
*P. megaterium* MOD5IV Promotes Arabidopsis thaliana Growth In Vitro. A) Promotion of *A. thaliana* primary root growth by the MOD5IV strain. B) Promotion of *A. thaliana* lateral root development induced by the MOD5IV strain. C) Growth of *A. thaliana* rosette induced by MOD5IV. a and b indicate significant differences estimated by Two-way ANOVA p < 0.05. Parametricity was estimated by Kolmogorov-Smirnov Test of Normality. N = 9. D) Representative images of *A. thaliana* plants exposed to MOD5IV and not exposed (Control) in a 15-day co-culture experiment.

### MOD5IV genomic features

To gain deeper insights regarding the genetic properties of the MOD5IV strain, we performed complete genome sequencing using Illumina and Nanopore sequencing platforms. 2,321,552,218 bp of raw Illumina data were obtained, corresponding to 15,374,518 100-bp reads. No quality filtering was required since the FasQC analysis indicated no overrepresented sequences and a quality Phred score above 30 throughout all the reads. The Nanopore platform generated 575,784,902 bp (56,256 reads, N50=32,731 bp). The complete genome was assembled using a hybrid approach, taking advantage of the higher-accuracy short reads from the Illumina platform and the significantly longer Nanopore reads.

The MOD5IV genome showed a total size of 5.69 Mb with an average GC content of 37.73%, consisting of a 5.25 Mbp circular chromosome and 9 plasmids designated pMOD5IV1 to pMOD5IV9 (Table 1, Figure 3A). Taxonomic classification according to the Genome Taxonomy Database confirmed this strain belonging to the *Priestia megaterium* species. A schematic representation of the replicons generated with the CG View server (Wang 2020; Stothard 2005) is provided in Figure 3A. Genome annotation showed 6,286 protein-coding genes, 75 repeat regions, and 189 RNA genes divided into 143 tRNAs and 46 rRNA (Table 1). The general features of the *P. megaterium* MOD5IV genome were compared with three genomes of *P. megaterium* strains with different numbers of plasmids (NTC-2, DMS319, KF18 and NBRC15305 = ATCC14581) and one *P. flexa* strain genome (KLBMP 4941) (Table 1). Homologous genome alignment drafts (Figure 3B) (Alikhan 2011) showed that the MOD5IV chromosome shared the general *Priestia megaterium* genome topology, except for two regions centered at 3 Mbp and 4.6 Mbp, which apparently correspond to insertions due to differences in the GC content. These insertions carried some open reading frames related to DNA repair and protein translation, such as Vsr-endonuclease and Aspartate-tRNA-synthetase. Also, the regions between 1.6 and 1.9 Mbp, around 3 Mbp, and between 3.6 and 3.8 Mbp showed a marked anomaly in the GC content, corresponding to divergent regions among the four strains compared. In these regions, it was possible to find genes related to sporulation, metal resistance, multidrug resistance, transcriptional regulators, chaperones, and DNA repair. Phage genes were detected only in the region around the coordinate 3.0 Mbp.

**Figure 3.**
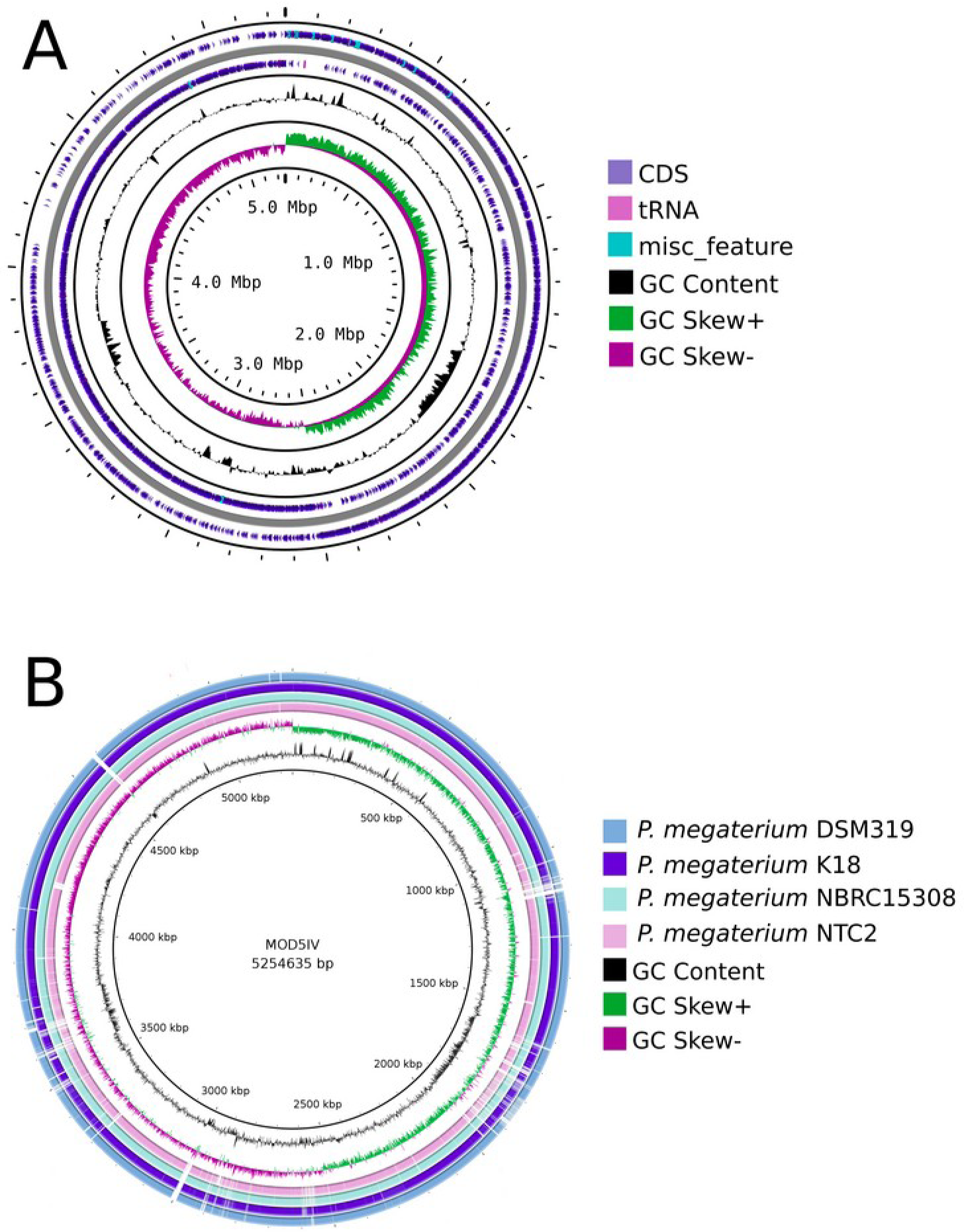
Genome map of *P. megaterium* MOD5IV and comparison with other P. megaterium genomes. A) Chromosome map prepared using Proksee software. Circles from outside to inside show the position of protein-coding sequences (blue), tRNA genes (pink), and rRNA genes (cyan) on the positive (circle 1) and negative (circle 2) strands. Circles 3 and 4 show plots of GC content and GC skew as deviations from the average for the entire sequence. B) BRIG output image. The draft of the P. megaterium MOD5IV genome aligned against other P. megaterium genomes (DM319, K18, NBRC15305 = ATCC 14581, NTC2). GC content and skew are also shown.

**Table 1.** General genome features of *P. megaterium* MOD5IV compared with other five *Priestia* strains.

| Strains | **P.**<br>**megaterium**<br>**MOD5IV** | **P.**<br>**megaterium**<br>**NCT-2** | **P.**<br>**megaterium**<br>**DSM 319** | **P.**<br>**megaterium**<br>**KF18** | **P.**<br>**megaterium**<br>**NBRC**<br>**15308** | **P. flexa**<br>**KLBMP**<br>**4941** |
| --- | --- | --- | --- | --- | --- | --- |
| **Genome size (Mb)** | 5.7 | 5.9 | 5.1 | 6.0 | 5.7 | 4.1 |
| **Chromosome size (Mb)** | 5.3 | 5.2 | 5.1 | 5.2 | 5.3 | 3.96 |
| **G+C content (%)** | 37.7 | 37.8 | 38.1 | 37.6 | 38.0 | 38.0 |
| **Chromosomal G+C content (%)** | 38.0 | 38.2 | 38.1 | 38.1 | 38.0 | 38.1 |
| **Gene number** | 6,211 | 6,134 | 5,337 | 6,294 | 6,003 | 4,332 |
| **Coding sequence number** | 6,022 | 5,805 | 4,994 | 5,979 | 5,754 | 3,971 |
| **RNA gene number** | 189 | 203 | 153 | 217 | 163 | 143 |
| **rRNA genes** | 46 | 53 | 33 | 46 | 41 | 36 |
| **tRNA gene number** | 243 | 142 | 114 | 163 | 122 | 102 |
| **Plasmid number** | 9 | 10 | 0 | 9 | 6 | 2 |

### MOD5IV genome functional annotation

To obtain better results in genomic annotation, the analysis was complemented using two different databases, RAST and KEGG (Shrestha 2022). The RAST platform annotated and functionally categorized 2,068 from a total of 5,722 coding sequences present in the MOD5IV genome. Of these, 2,025 were found in the chromosome, with the remainder distributed among the different plasmids. A total of 1,361 chromosomal genes (25%) were assigned to 334 subsystems by RAST. The top five categories in strain MOD5IV were “Amino Acids and Derivatives” (414), “Carbohydrates” (318), “Cofactors, Vitamins, Prosthetic Groups, Pigments” (159), “Protein Metabolism” (243), and “Nucleosides and Nucleotides” (108) (Figure 4A). No matches were found for the “Photosynthesis” and “Phages, Prophages, Transposable Elements, Plasmids” categories. Meanwhile, the KEGG platform identified 6,022 genes, with 2,915 genes annotated and classified into KEGG categories. The top five categories were “Protein Families: Genetic Information Processing” (381), “Carbohydrate Metabolism” (343), “Protein Families: Signaling and Cellular Processes” (324), “Environmental Information Processing” (227), and “Unclassified: Metabolism” (199) (Figure 4B).

**Figure 4.**
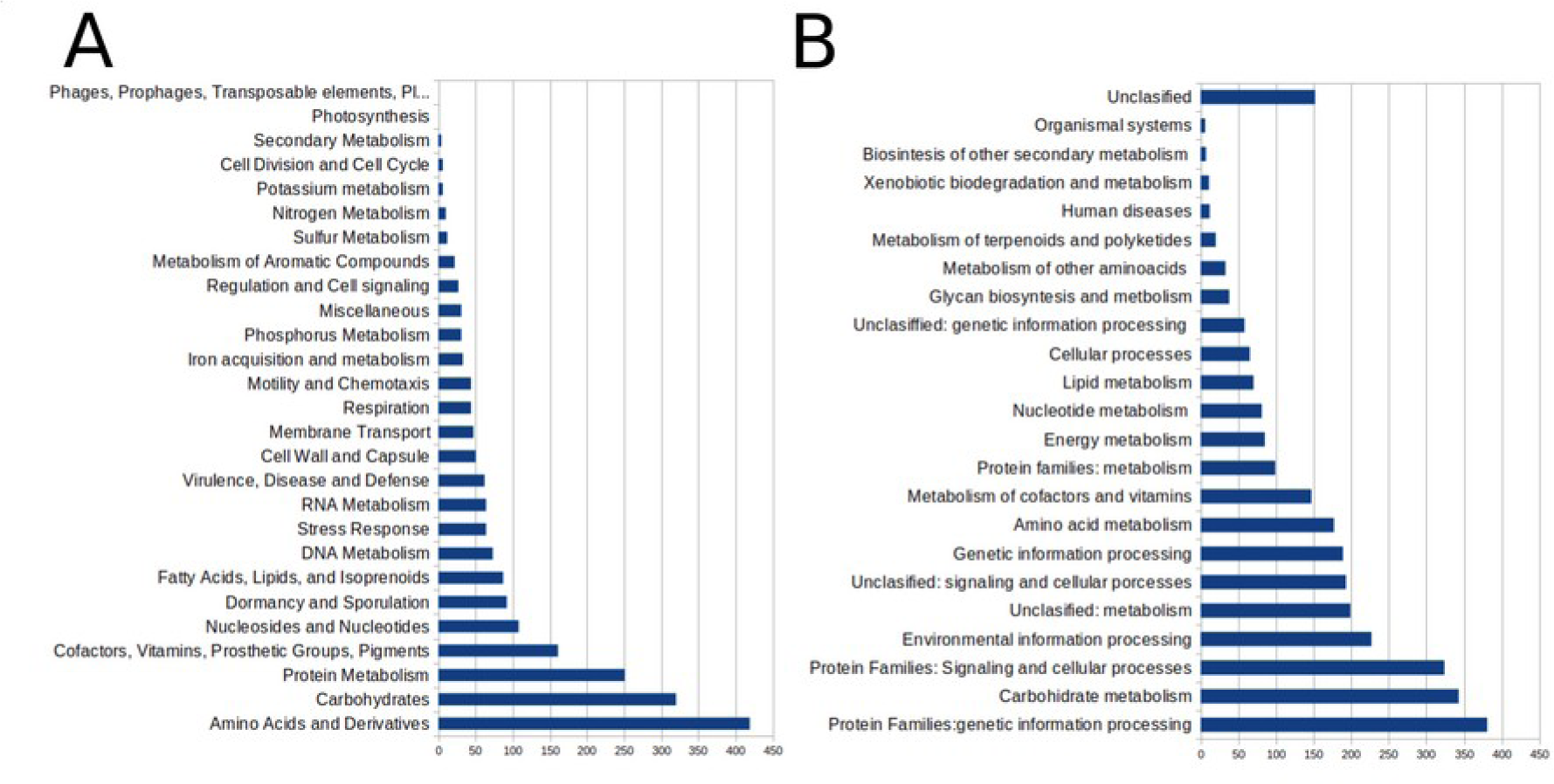
Predicted functions encoded in the *P. megaterium* MOD5IV genome. A) Statistical chart of the RAST subsystem category distribution. The y-axis represents the classification of different functional genes in the RAST subsystem classification. B) Statistical chart of the KEGG annotation classes. The y-axis represents the classification of different functional genes in the KEGG database.

The proteome of MOD5IV was extracted from the RAST server and submitted to the FusionBD server (https://services.bromberglab.org/fusiondb/) to search for similarities with related strains. The MOD5IV proteome contained 5449 proteins, which were mapped to 3780 functions. Five hundred seventy-three proteins could not be mapped to any function in the BD database. The most similar proteome found in the database was *P. megaterium* DSM319 (TAXID 592022) with 81% similarity and 3511 shared functions.

Core and accessory genomes were determined using the Spine/AGEnt/ClustAGE tool (http://spineagent.fsm.northwestern.edu/index_age.html) and 21 *Priestia megaterium* reference strains (DSM319, YC4-R4, KF18, 207, NCT-2, KNU-01, 5-3, S188, S2, FDU301, BIM B-1314D, NBRC15305 = ATCC14581, BHS1, IGA-FME-1, CDC 2008724142, H2, 2020WEIHUA_L, BP01R2, CK7, F20, and KF18) available in BioProject database with an assembly level of complete genome. Approximately 25.5% of the MOD5IV genome was accessory. The core genome (4,239,675 bp) consisted of 4,581 coding sequences with a GC content of 38.7%. Meanwhile, the accessory genome had 1,783 coding sequences and a GC content of 34.9%. The differences in GC content between the core and accessory genomes could indicate horizontal gene transfer. RAST annotation of the accessory genome revealed that the principal subsystem categories corresponded to “Amino Acids and Derivatives,” “Stress Response,” and “Cell Wall and Capsule.” This differed from the core genome, where the principal categories were “Amino Acids and Derivatives,” “Carbohydrates,” and “Protein Metabolism” (Figure 5). The presence of genes classified into the “Virulence, Disease, and Defense” subsystem also increased considerably in the accessory genome of MOD5IV, including genes related to copper homeostasis, cobalt-zinc-cadmium resistance, and antibiotic resistance. Meanwhile, in the “Stress Response” subsystem, the accessory genome of MOD5IV contained genes related to osmotic stress (choline and betaine uptake), cold shock, and bacterial hemoglobins. A total of 429 coding sequences were present in the nine plasmids of MOD5IV (Table 2). In the best case, RAST annotation covered no more than 14% of the sequences. Among the annotated sequences, we found genes related to isoprenoid biosynthesis, sugar ABC transporters, oxidative stress (catalase), osmotic stress (glycine/betaine transporters or choline-binding protein A), and phage capsid genes.

**Figure 5.**
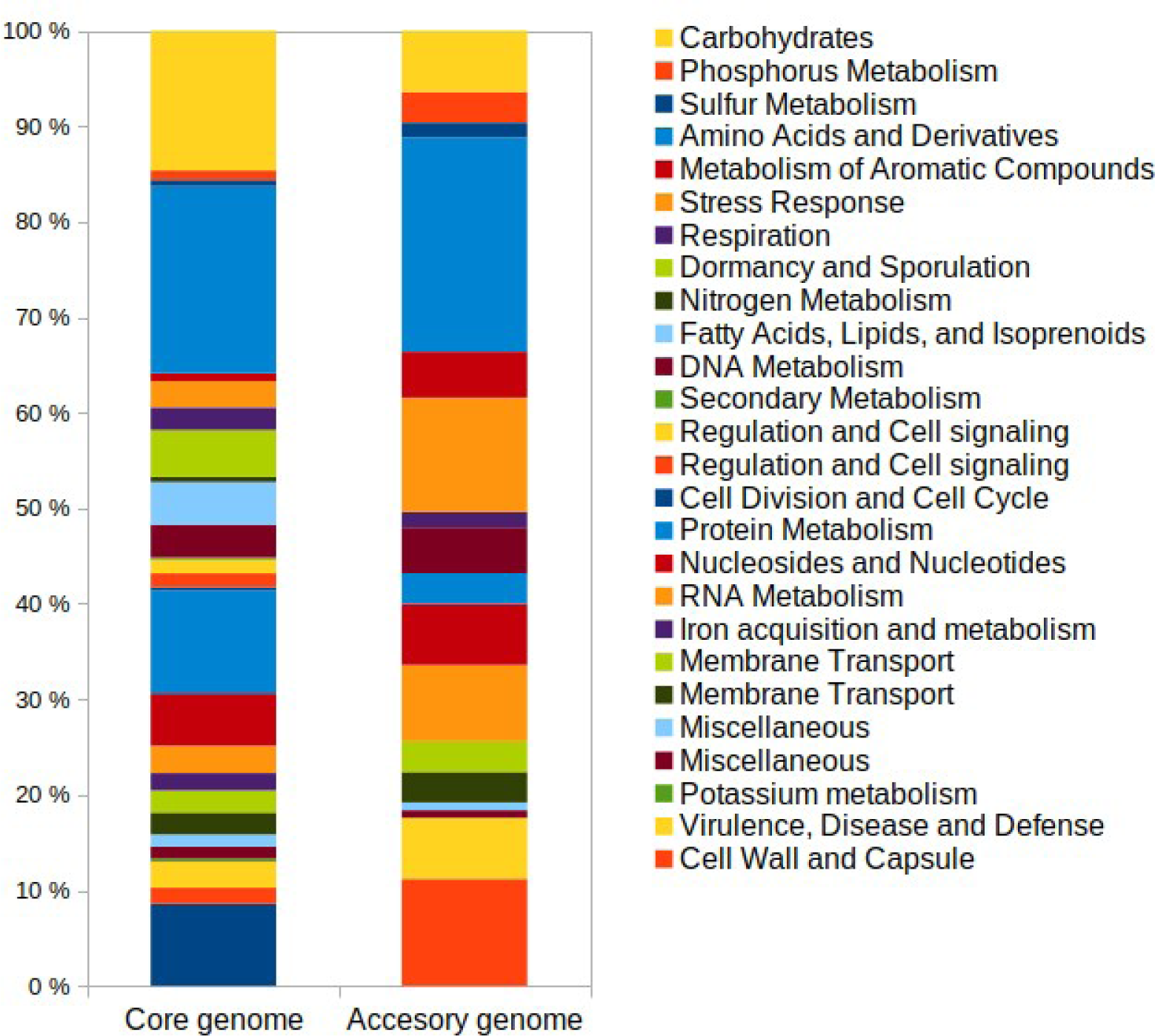
*P. megaterium* MOD5IV Accessory Genome Analysis. RAST subsystem classification of the MOD5IV core and accessory genome determined by Spine/AGEnt. Data is presented as a percentage of the total genes in the core and accessory genome.

**Table 2.** General features of the chromosome and plasmids present in *P. megaterium* MOD5IV genom.

|  | **Chrom**<br>**osome** | **pMOD**<br>**5IV**<br>**1** | **pMOD**<br>**5IV**<br>**2** | **pMOD**<br>**5IV**<br>**3** | **pMOD**<br>**5IV**<br>**4** | **pMOD**<br>**5IV**<br>**5** | **pMOD**<br>**5IV**<br>**6** | **pMOD**<br>**5IV**<br>**7** | **pMOD**<br>**5IV**<br>**8** | **pMOD**<br>**5IV**<br>**9** |
| --- | --- | --- | --- | --- | --- | --- | --- | --- | --- | --- |
| **Size (pb)** | 5,254,635 | 125,337 | 112,536 | 69,915 | 47,78 | 42,938 | 10,013 | 9,405 | 8,643 | 8,518 |
| **GC**<br>**Content** | 38.0 | 34.1 | 34.5 | 31.7 | 32.0 | 37.3 | 34.8 | 34.4 | 33.8 | 34.5 |
| **Number of**<br>**RAST**<br>**Subsystem**<br>**s** | 334 | 9 | 6 | 6 | 0 | 0 | 0 | 0 | 0 | 1 |
| **Number of**<br>**Coding**<br>**Sequences** | 5.554 | 140 | 62 | 84 | 53 | 42 | 14 | 10 | 13 | 11 |
| **Number of**<br>**RNAs** | 168 | 0 | 0 | 0 | 0 | 20 | 0 | 0 | 0 | 0 |

### Plant growth promotion and resistance genes

As expected, the MOD5IV genome contained several genes related to both direct and indirect mechanisms of plant growth promotion (Backer 2018). Regarding direct mechanisms, MOD5IV included ten genes related to PO₄ solubilization and uptake, such as *phoU*, *phoP*, *phoR*, *phoB*, and *phoH*; two genes related to potassium uptake, *kefA* and the large-conductance mechanosensitive K channel; 21 genes related to siderophore production and Fe assimilation; and 12 genes related to nitrogen assimilation, including *norD*, *norQ*, nitrate reductase, and glutamine synthase, among others. Concerning indirect plant growth-promoting mechanisms, MOD5IV housed 10 genes related to auxin production (indole-acetic acid, IAA), primarily belonging to the indole-3-pyruvic acid and tryptamine synthesis pathways (Supplementary Table 2).

The MOD5IV genome showed a variety of metal resistance genes distributed principally into the chromosome. These include arsenic resistance genes grouped into two putative operons: one included *arsC*, *arsB*, and *arsR*, while the other contained only *arsR* and *arsB*. Additionally, copper resistance genes were grouped into three putative operons: one operon encoded a copper-translocating P-type ATPase, *copD*, and *ycnl*; another contained *copC*, *copD*, and *cutC*; and the third encoded a copper-translocating P-type ATPase, *copZ*, and *csoR*. Cadmium resistance genes *cadA* and *cadC* were grouped into one putative operon. Three mercury-sensitive transcriptional regulator genes (*merR)* not organized into any *mer* operon were present in the genome. Resistance genes for other metals such as lead, cobalt, and zinc were also present and organized in an operon containing the *P-type Cd^2+/^Zn^2+^ transporter*, the transcriptional repressor *czrA*, and the cobalt/zinc/cadmium resistance protein *czcD* (Supplementary Table 2).

Using the Resistance Gene Identifier (RGI) tool (Alcock 2023), we further characterized the MOD5IV resistome, identifying 12 probable antibiotic resistance genes, which include six genes for glycopeptide antibiotic resistance (*vanY/I/B/W/G/T*), one for tetracycline resistance (*tetQ*), one for cephalosporin resistance *(bcIII*), one for fosfomycin resistance (*fosB*) and three small multidrug resistance efflux pump (*qacG/J*).

### MOD5IV metal resistance and phytoremediation potential

Metal exposure experiments were performed to establish the resistance range of MOD5IV. The strain was exposed to increasing concentrations of common contaminating metals diluted in TSB medium, and growth kinetics were monitored by measuring the absorbance at 600 nm. Figure 6 summarizes the resistance level of the MOD5IV strain, which exhibited high tolerance to multiple metals (and metalloids), including arsenic, chromium, lead, and copper. Conversely, mercury and cadmium induced toxic effects at low concentrations.

**Figure 6.**
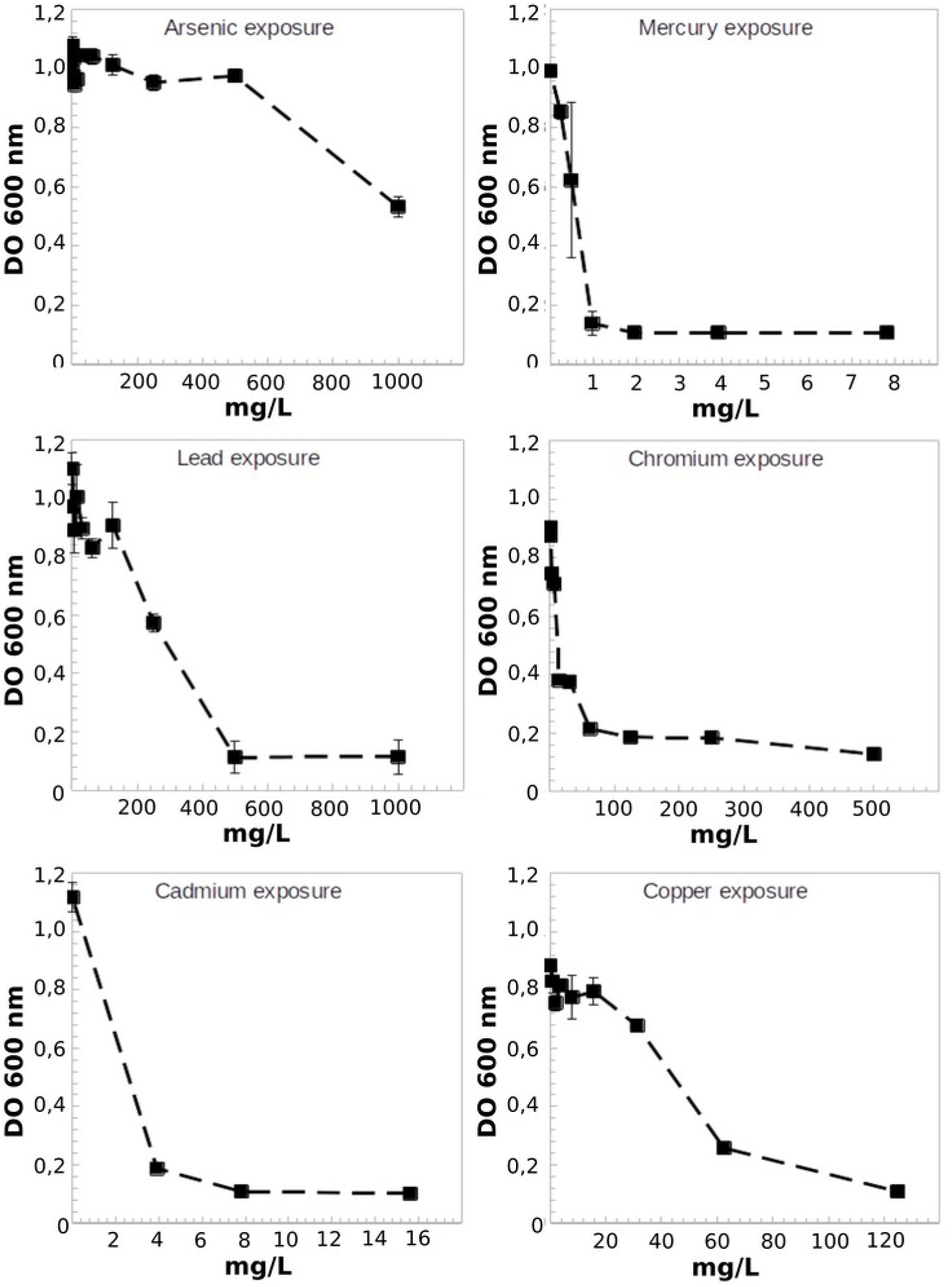
*P. megaterium* MOD5IV Metal Resistance. Maximum growth of MOD5IV in TSB medium supplemented with increasing concentrations of metals (as indicated in each figure), measured by absorbance at 600 nm. Error bars indicate the standard deviation of the average of three independent experiments. MIC: As = >1000 mg/L; Hg = 2 mg/L; Pb = >500 mg/L; Cr = 500 mg/L; Cd = 8 mg/L; Cu = 120 mg/L.

MOD5IV was initially isolated from the rhizosphere of a *C. spinosa* specimen. Accordingly, the same plant species was used to test the capabilities of MOD5IV to promote plant growth and improve phytoremediation potential in conditions of high metal concentrations in the substrate. A co-culture experiment was conducted in which *C. spinosa* seedlings were grown for 60 days in tailings material amended with organic substrate and treated with MOD5IV. After the culture period, the seedlings were extracted, and growth parameters were measured. As could be seen, seedlings inoculated with MOD5IV showed a significant increase in growth parameters in comparison with non-inoculated seedlings, including the length of roots and aerial parts (shoots + leaves) (Figure 7A) and dry weight (Figure 7B) parameters. Additionally, an increase in Cu uptake was recorded (Figure 7C), corresponding to an increase in Cu concentration of approximately 63% in the aerial parts and 29% in the roots. Although *C. spinosa* appears to be very tolerant to conditions of high metal concentrations in the substrate, it is not a hyperaccumulating plant, as the Cu concentration in the roots and aerial parts showed an accumulation of only 3% and 0.6%, respectively, of the copper present in the substrate, which registers 1,974 mg(Cu)/kg.

**Figure 7.**
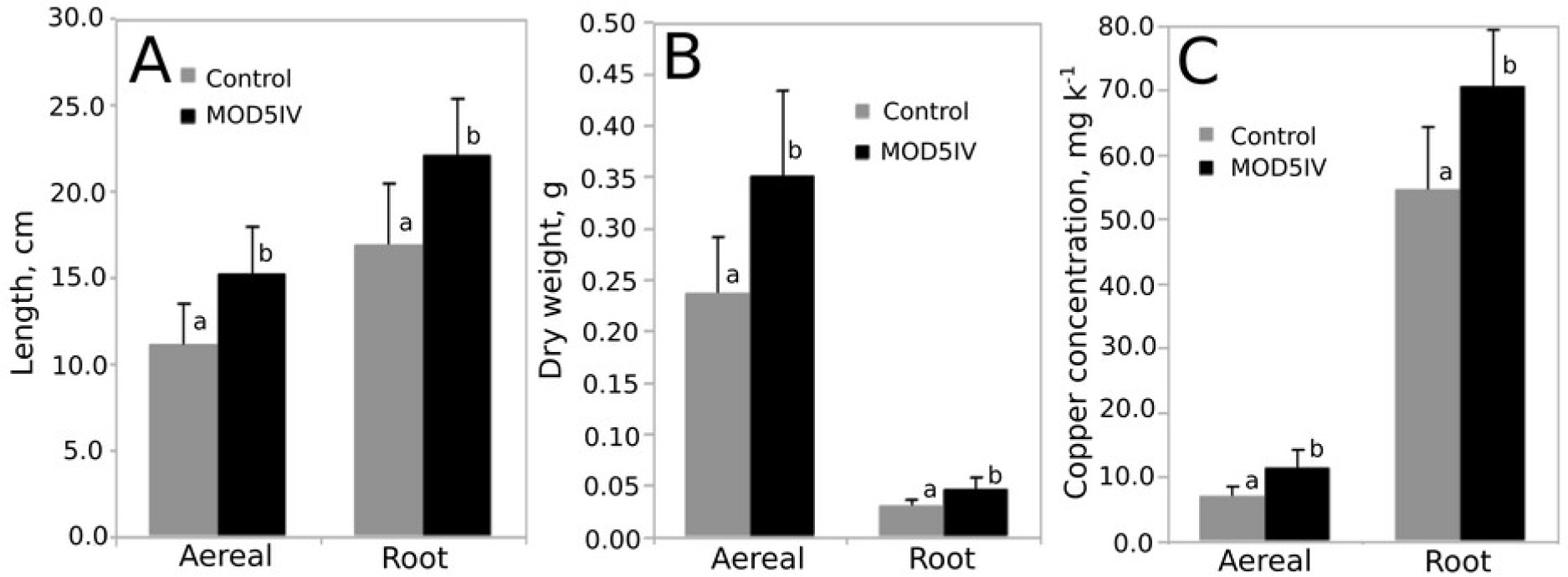
*P. megaterium* MOD5IV Increases Growth and Copper Uptake in Caesalpinia spinosa. Two-month-old C. spinosa plants were grown in mine tailings amended with organic substrate, with or without inoculation with P. megaterium MOD5IV. A) aerial (shoot + leafs) and root length of C. spinosa. B) Dry weight of C. spinosa aerial organs and roots. C) Copper accumulation in aerial organs and roots of C. spinosa. a and b indicate significant differences estimated using Two-way ANOVA (p < 0.05). Parametricity was estimated by the Kolmogorov-Smirnov Test of Normality.

## DISCUSSION

Phytoremediation efforts of metal-contaminated soils largely benefit from bacterial strains promoting the growth of plants performing the bioaccumulation process. These strains must have two concomitant features that do not always come together: the capacity to resist the highly toxic metals present, and the metabolic capacities leading to (ideally several) plant growth-promoting effects. As stated above, the MOD5IV genome contains a complex array of metal resistance genes and operons, conferring resistance to multiple metals, as has been previously reported for other *P. megaterium* strains. For example, the strain HgT21 contains two complete type II *mer* operons in its genome and tolerates about 6.8 mg/L of Hg (Guzmán-Moreno 2022). MOD5IV contains the transcriptional regulator *merR* gene repeated three times in its genome but lacks the rest of the *mer* operon elements. Despite this, MOD5IV could resist 2 mg/L of Hg^2+^ in solution. The HgT21 strain also contains the *ars* operon and tolerates high concentrations of arsenic (5,697.3 mg/L As(V)) in the form of Na_3_AsO_4_, whereas MOD5IV tolerated over 1,000 mg/L of arsenite (As(III)) (as As_2_O_3_). This high tolerance correlates with the presence of two *ars* operons in its genome. Higher concentrations of As(III) could not be tested due to its low solubility in the culture medium. MOD5IV also resisted the presence of a high concentration of copper in the culture medium (MIC 125 mg(Cu)/L) at a similar range as the *P. megaterium* HgT21 strain (190 mg(Cu)/L). This resistance could be linked to the presence of three putative copper operons in its genome, including one *csoR-copZA* system that has been described to confer resistance to high levels of copper (Efe 2020, Smaldone 2007).

Regarding cadmium resistance, MOD5IV contained one *cad* operon and other operons conferring resistance to lead, zinc, cadmium, and cobalt, resulting in a resistance of 8 mg/L of Cd in solution, which is slightly greater than the previously reported Cd-resistant *P. megaterium* BM18 strain that has demonstrated resistance of over 4.5 mg/L of Cd (Wu 2019). MOD5IV also showed high resistance to Pb with a MIC of 500 mg/L, considerably more than other *P. megaterium* isolates, which have shown a resistance of 20.7 mg/L of Pb in solution (Roane 1999). However, strains of the genus *Bacillus* have shown resistance to similar amounts of Pb (Abdelkrim 2018). Finally, MOD5IV showed a MIC of 250 mg/L of Cr, despite the absence of specific Cr resistance genes, such as the *chr* operon. Other *P. megaterium* isolates have resisted 150 mg/L of Cr(VI) (Srinath 2002). Notably, MOD5IV presents resistance to a greater number of different metals than other reported *P. megaterium* strains.

Several studies have demonstrated the ability of metal-resistant PGPB strains to increase plant growth and phytoremediation potential in metal-contaminated soils (Bhatt 2020, Chakraborty 2006, Tirry 2021, Tang 2020, Ustiatik 2021, Ullah 2022, Kong 2015, Gao 2010, Ma 2011), including some belonging to *P. megaterium* taxa as the strain HgT21 (Guzmán-Moreno 2022) and M18-2 (Wu 2019). To our knowledge, MOD5IV is the first multi-metal resistant and PGPB *P. megaterium* strain reported with the capacity to increase the phytoremediation potential in experiments using copper mining waste in the substrate. Plant growth-promoting bacteria can act by reducing the toxic effects of metals or by producing phytohormones (Wu 2019). Depending on the PGP traits, different bacteria play a key role in promoting plant development besides reducing the toxicity or damage to plants exposed to stress from different heavy metals. Additionally, PGPB strains produce various metabolic compounds such as organic acids, siderophores, EPS, and biosurfactants, which can alter the redox state, chelation, precipitation, and immobilization of the metals in the rhizospheric environment, thereby enhancing the plant phytoremediation efficiency (Manoj 2020, Alves 2022).

Biosurfactants, as the lipopeptide surfactin, have shown to play an important role in metal bioremediation (Mishra 2021) and copper bioleaching (Rozas 2019). Upon searching the MOD5IV genome, we found no genetic determinants for the surfactin synthesis pathway, despite of its synthesis has been reported for another *P. megaterium* strain (Cui 2023). Considering the above, it is likely that copper resistance genes may be crucial in conferring phytoremediation efficiency to *C. spinosa* plants. Specifically, the *copD* and *copC* genes, which are involved in the import of copper into the cytoplasm (Lawton 2016), and the cytoplasmic copper-binding protein CutC (Latorre 2011) reduce copper toxicity and facilitate its safe transport within the cell (Rensing 2003). Additionally, the phosphate solubilization ability of the tested bacterial strain likely plays an important role in enhancing plant uptake of soil minerals and increasing the uptake of metals in contaminated soils through the secretion of organic acids (Manoj 2020, Ma 2011). Bacteria such as MOD5IV and other metal-tolerant rhizobacteria selected by plants, are crucial for metal detoxification in metalliferous soils. They help plants endure metal toxicity and improve metal uptake. Thus, the efficiency of phytoextraction can be enhanced by promoting both plant growth and metal bioavailability in the soil (Ashraf 2019).

Summarizing, The strain *P. megaterium* MOD5IV exhibited outstanding genetic and phenotypic features, which related plant growth promotion and metal resistance with the increase in the phytoremediation potential of the desert woody plant *C. spinosa*. These results provide valuable genomic and experimental resources for further studies and applications of *P. megaterium* MOD5IV in bioremediation processes, particularly in heavy metal-contaminated soils. Finally, the identification of the *P. megaterium* MOD5IV strain isolated from metal-contaminated soils in the arid regions of South America, increases the evidence of the Atacama desert as a source of microbiological solutions for climate adaptation and environmental remediation (Castro-Severyn 2024, Astorga-Eló 2021, Maza 2019).

## ACKNOWLEDGMENTS

We thank Duna Soluciones Biotecnológicas SpA. for providing materials and samples used in this study.

