## Supplementary Table 1, Suplementary Figure 1 and Supplementary Table 2 for "Genomic and functional insights on *Priestia megaterium* MOD5IV: Enhancing Metal Phytoremediation Potential in Arid Environments"

**Supplementary Table 1.** Concentration of elements and compounds present in the tailing material from where the MOD5IV strain was isolated. A soil sample was collected during bacterial isolation, from a 1 m × 1 m area to a depth of 15 cm using a shovel. The amount of soil in each sample was then reduced using the quartering method (ASTM International, 2018) to a representative 1000 g of the sample. The sample was analyzed using X-ray fluorescence (XRF) with an S1 Titan 600 Handheld XRF Analyzer (Bruker). Error values indicate standard deviations between 5 measurements.

| Element / compound | Concentration (mg k-1) | Error |
|---|---|---|
| **SiO2** | 381.639 | 2.987 |
| **Al2O3** | 109.330 | 2.946 |
| **Fe2O3** | 82.281 | 310 |
| **K2O** | 25.068 | 166 |
| **MgO** | 18.366 | 12.704 |
| **S** | 17.335 | 236 |
| **CaO** | 5.643 | 97 |
| **TiO2** | 5.056 | 78 |
| **Zn** | 3.216 | 37 |
| **Cu** | 3.140 | 38 |
| **MnO** | 1.928 | 67 |
| **P2O5** | 1.475 | 379 |
| **Cl** | 983 | 150 |
| **Ba** | 810 | 197 |
| **Ce** | 435 | 98 |
| **Pb** | 309 | 35 |
| **Zr** | 288 | 10 |
| **V** | 220 | 24 |
| **Sr** | 216 | 8 |
| **Mo** | 184 | 15 |
| **Rb** | 178 | 11 |
| **Co** | 97 | 50 |
| **W** | 81 | 76 |
| **As** | 63 | 45 |
| **Y** | 60 | 4 |

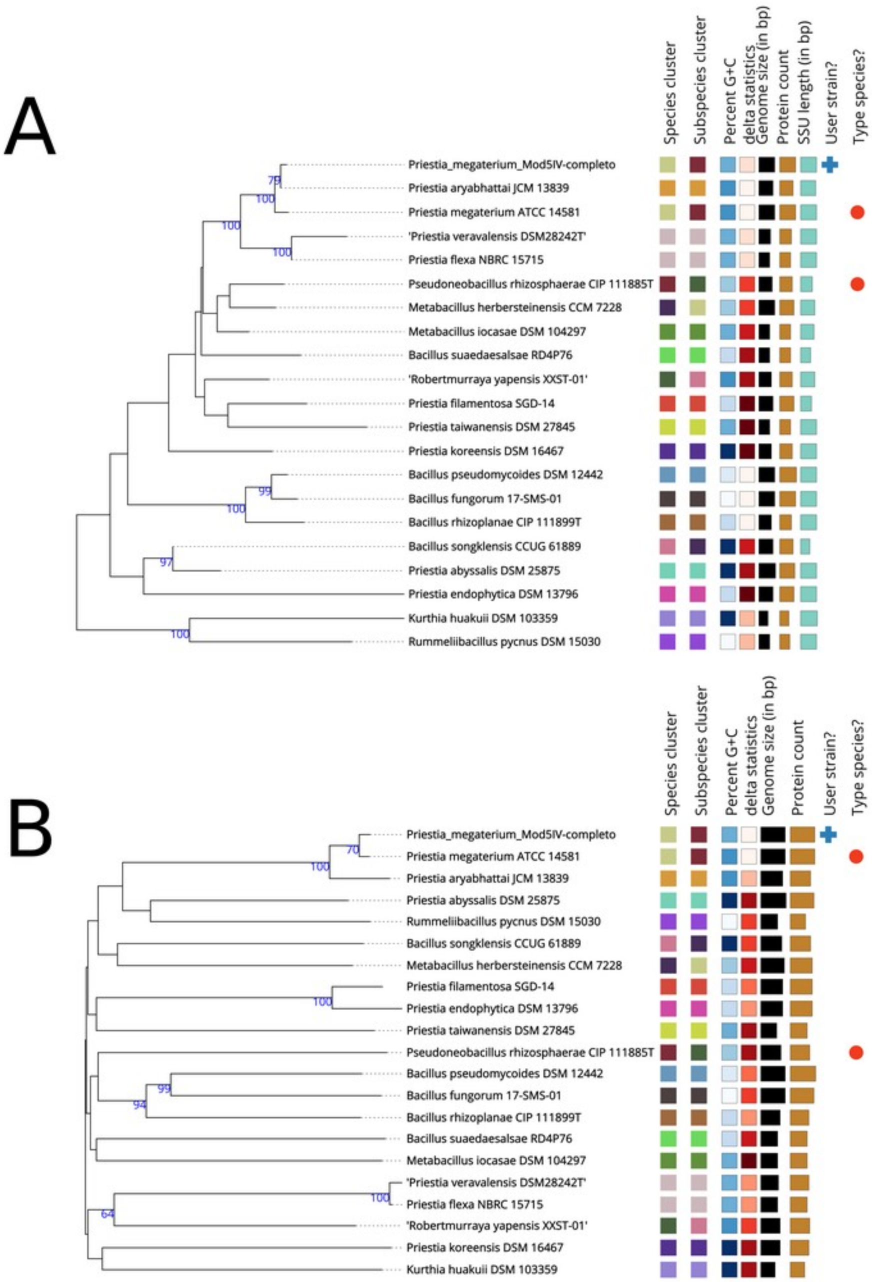


**Supplementary Figure 1.** Phylogenetic analysis of MOD5IV elaborated in TYGS (Meier-Kolthoff et al 2022) A) 16S rDNA gene sequence-based Genome BLAST Distance Phylogeny tree (GBDP) and B) whole-genome sequence-based GBDP tree.

**Supplementary Table 2.** Summary of Gene Functions Related to Plant Growth Promotion and Resistance to Metals and Metalloids in the *P. megaterium* MOD5IV Genome.

| | **Gene function** |
|---|---|
| **N.º** | **Fe metabolism and uptake** |
| **1** | Sdab Siderophore biosynthesis diaminobutyrate-2-oxoglutarate aminotransferase |
| **2** | Sdab Siderophore biosynthesis L-2,4-diaminobutyrate decarboxylase |
| **3** | SSIAc Siderophore synthetase large component, acetyltransferase |
| **4** | SssAc Siderophore synthetase small component, acetyltransferase |
| **5** | SSLc Siderophore synthetase component, ligase |
| **6** | SbsM Siderophore biosynthesis protein, monooxygenase |
| **7** | Stra Siderophore transport protein |
| **8** | ABCp ABC-type Fe3+-siderophore transport system, permease component |
| **9** | ABCatp ABC-type Fe3+-siderophore transport system, ATPase component |
| **10** | ABCsb ABC-type Fe3+-siderophore transport system, periplasmic iron-binding component |
| **11** | ABCp2 ABC-type Fe3+-siderophore transport system, permease 2 component |
| **12** | X-ABC3 ABC Uncharacterized iron compound ABC uptake transporter, ATP-binding protein |
| **13** | Fe-ABC1 Iron compound ABC uptake transporter substrate-binding protein |
| **14** | Fe-ABC2 Iron compound ABC uptake transporter permease protein |
| **15** | HtsABC Heme ABC type transporter HtsABC, heme-binding protein |
| **16** | HrtAB Sensor histidine kinase colocalized with HrtAB transporter |
| **17** | X-ABC Uncharacterized iron compound ABC uptake transporter, ATP-binding protein. |
| **18** | Iron compound ABC uptake transporter permease protein |
| **19** | ZnH Zn-dependent hydrolase YycJ/WalJ |
| **20** | Two-component response regulator SA14-24 |
| **21** | Two-component sensor kinase SA14-24 |
| **N.º** | **Phosphate metabolism and uptake** |
| **1** | PhoU Phosphate transport system regulatory protein |
| **2** | PhoP Alkaline phosphatase synthesis transcriptional regulatory protein |
| **3** | PhoR Phosphate regulon sensor protein (SphS) |
| **4** | PhoB Phosphate regulon transcriptional regulatory protein (SphR) |
| **5** | Polyphosphate kinase |
| **6** | Manganese-dependent inorganic pyrophosphatase |
| **7** | PhoH Predicted ATPase related to phosphate starvation-inducible protein |
| **8** | Alkaline phosphatase |
| **9** | Probable low-affinity inorganic phosphate transporter |
| **10** | Exopolyphosphatase |
| **N.º** | **Potassium metabolism and uptake** |
| **1** | Large-conductance mechanosensitive channel. |
| **2** | Potassium efflux system KefA protein |
| **N.º** | **Auxin biosyntesis** |
| **1** | Anthranilate phosphoribosyltransferase |
| **2** | Phosphoribosylanthranilate isomerase |
| **3** | Tryptophan synthase alpha chain |
| **4** | Tryptophan synthase beta chain |
| **5** | Amino transferase |
| **6** | Indolepyruvate decarboxylase |
| **7** | IAAld dehydrogenase |
| **8** | Trp decarboxylase |
| **9** | Tryptophan acetyltransferase |
| **10** | Nitrilase |

| N.º | Nitrogen metabolism and uptake |
|---|---|
| 1 | NsrR Nitrite-sensitive transcriptional repressor |
| 2 | Glutamine synthetase type I |
| 3 | Glutamate synthase [NADPH] small chain |
| 4 | Ferredoxin-dependent glutamate synthase |
| 5 | Nitrogen regulatory protein P-II |
| 6 | Ammonium transporter |
| 7 | Assimilatory nitrate reductase large subunit |
| 8 | Nitrite reductase [NAD(P)H] large subunit |
| 9 | Nitrate/nitrite transporter NarT |
| 10 | NorD, Nitric oxide reductase activation protein |
| 11 | NorQ, Nitric oxide reductase activation protein |
| 12 | Oxygen-insensitive NAD(P)H nitroreductase |
| **N.º** | **Metal resistance** |
| 1 | MerR Transcriptional regulator, MerR family |
| 2 | ArsR Arsenical resistance operon repressor |
| 3 | ArsC Arsenate reductase thioredoxin-coupled, LMWP family |
| 6 | ArsB Arsenite/antimonite:H+ antiporter |
| 7 | Lead, cadmium, zinc and mercury transporting ATPase |
| 8 | Multidrug resistance transporter, Bcr/CflA family |
| 9 | Copper resistance protein CopC |
| 10 | Copper resistance protein CopD |
| 11 | Cobalt-zinc-cadmium resistance protein |
| 12 | Cadmium-transporting ATPase |
| 13 | Response regulator of zinc sigma-54-dependent two-component system |
| 14 | Cytoplasmic copper homeostasis protein CutC |
| 15 | CadA Cadmium-transporting ATPase |
| 16 | CadC Cadmium efflux system accessory protein |
